# Sex checking by zygosity distributions

**DOI:** 10.64898/2026.03.15.711924

**Authors:** Oscar Molina-Sedano, Daniel Mas Montserrat, Alexander G. Ioannidis

## Abstract

**Motivation:** In genomic and clinical studies, verifying concordance between self-reported and genotype-inferred sex is a crucial quality control step, since mismatches arising from mislabeling or aneuploidies can bias downstream analyses and affect diagnostic accuracy. Existing approaches typically require substantial auxiliary data, and often require manual threshold tuning. There remains a need for a streamlined, reference-free method that generalizes across different data modalities—including whole-genome, single-sample and array—without requiring additional files or parameter tuning.

**Results:** We present Zigo, a novel ML-based sex-checking method that operates solely on a standard VCF file, designed using X-chromosome genotype class distributions across sexes. Our model was trained on synthetic data incorporating standard demographic models and empirical recombination maps to ensure realistic genetic architecture and population structure. We simulate WGS, array, and single-sample files for broad applicability. Unlike traditional methods, we eliminate manual thresholding by distilling learned discriminative patterns into a single polynomial equation that determines genetic sex directly from normalized genotype counts. We validated Zigo on independent datasets, including 1000 Genomes, UK Biobank, and HGDP. Additional experiments assessed robustness under reduced variant availability through random SNP subsampling and allele-frequency filtering. Across all evaluations, the model achieved state-of-the-art accuracy, high time efficiency, and strong generalization, even with severely limited variant sets.

**Availability:** We release our sex-checking tool as an open-source command-line interface (CLI) under GitHub at https://github.com/AI-sandbox/zigo.

**Supplementary information:** Supplementary data are available in the Supplementary Material at the end of this article.

## 1 Introduction

Ensuring data integrity is a fundamental prerequisite in genomic research, where errors propagated from upstream processing can severely compromise downstream association analyses and clinical interpretations. Between quality control (QC) procedures, verifying the concordance between self-reported and genotype-inferred sex is a critical safeguard against sample misidentification and clerical errors [5, 15, 25]. Large-scale biobanking initiatives, such as the UK Biobank and the All of Us Research Program, implement rigorous multi-step QC pipelines where sex concordance checking serves as a primary filter to detect sample swaps and exclude individuals with sex chromosomal aneuploidies or discordant metadata prior to association testing [4, 26].

Despite its importance, computationally inferring genetic sex remains a challenge due to the diversity of sequencing technologies and data formats. Several established methods rely on read depth or alignment metrics from BAM/CRAM files to compare X and Y chromosome coverage ratios [17, 27]. While effective in many scenarios, these alignment-based approaches impose significant storage and require access to raw alignment data that may not be available in shared summary datasets. Furthermore, many algorithms explicitly require Y-chromosome data to function effectively [21], which limits their applicability in targeted panels or assays where the Y chromosome is not captured or is filtered out to reduce file size.

For genotype-only datasets (VCF/PLINK format), the industry standard relies on calculating the X-chromosome inbreeding coefficient (*F*) to assess heterozygosity, a method implemented in widely used tools like PLINK, Hail or peddy [10, 18, 20, 23]. Theoretically, this approach exploits the Hardy-Weinberg Equilibrium (HWE), expecting females (XX) to exhibit diploid heterozygosity rates (*F*≈ 0) and males (XY) to be effectively haploid (*F* ≈1). However, this metric is sensitive to the underlying population allele frequencies used for calculation. As highlighted by Backenroth and Carmi [6], deviations from expected genotype frequencies in sex-biased admixed populations can confound standard HWE-based tests, leading to ambiguous classification in the intermediate range (0.2 *< F <* 0.8) [6]. Consequently, these methods often require defining external reference panels for accurate frequency estimation or demand manual post-hoc tuning of decision thresholds to accommodate dataset-specific noise, steps that hinder automated, reproducible workflows. Probabilistic approaches, such as the Hidden Markov Models (HMM) employed by BCFtools [11], offer distinct advantages by modeling ploidy states directly. However, these methods typically estimate allele frequencies directly from the input data. In single-sample files, this internal estimation is insufficient without external population references, potentially leading to inaccurate inferences. In practice, these techniques require input files to be homogenized with respect to reference panels which can be difficult and time consuming. Homogenizing datasets and computing allele frequencies can be a time consuming task difficulting the adoption of current sex-checking techniques (PLINK, Hail, BCFtools). Therefore, there is still a need for methods that can effectively predict genetic sex on single-sample variant call files without requiring the merging with information from additional sequence individuals.

Here, we present Zigo, a novel reference-free method that infers genetic sex solely from X-chromosome variant counts. Our approach leverages a gradient boosting framework, subsequently distilled into a single, interpretable polynomial equation. By analyzing the distribution of normalized genotype class counts, this model captures discriminative patterns that transcend simple heterozygosity ratios. This formulation allows the method, trained on realistic synthetic data, to achieve state-of-the-art accuracy and generalization across whole-genome sequencing (WGS), genotyping arrays, and single-sample files without requiring external reference panels, Y-chromosome data, or manual threshold tuning. Our approach has been validated on independent, real-data benchmarks including the 1000 Genomes Project, Human Genome Diversity Project (HGDP), and UK Biobank datasets. As visualized in Figure 1, our distilled polynomial equation establishes a robust, universal decision boundary that accurately segregates sexes across these heterogeneous data modalities within the unified geometric space of genotype frequencies.

**Figure 1:**
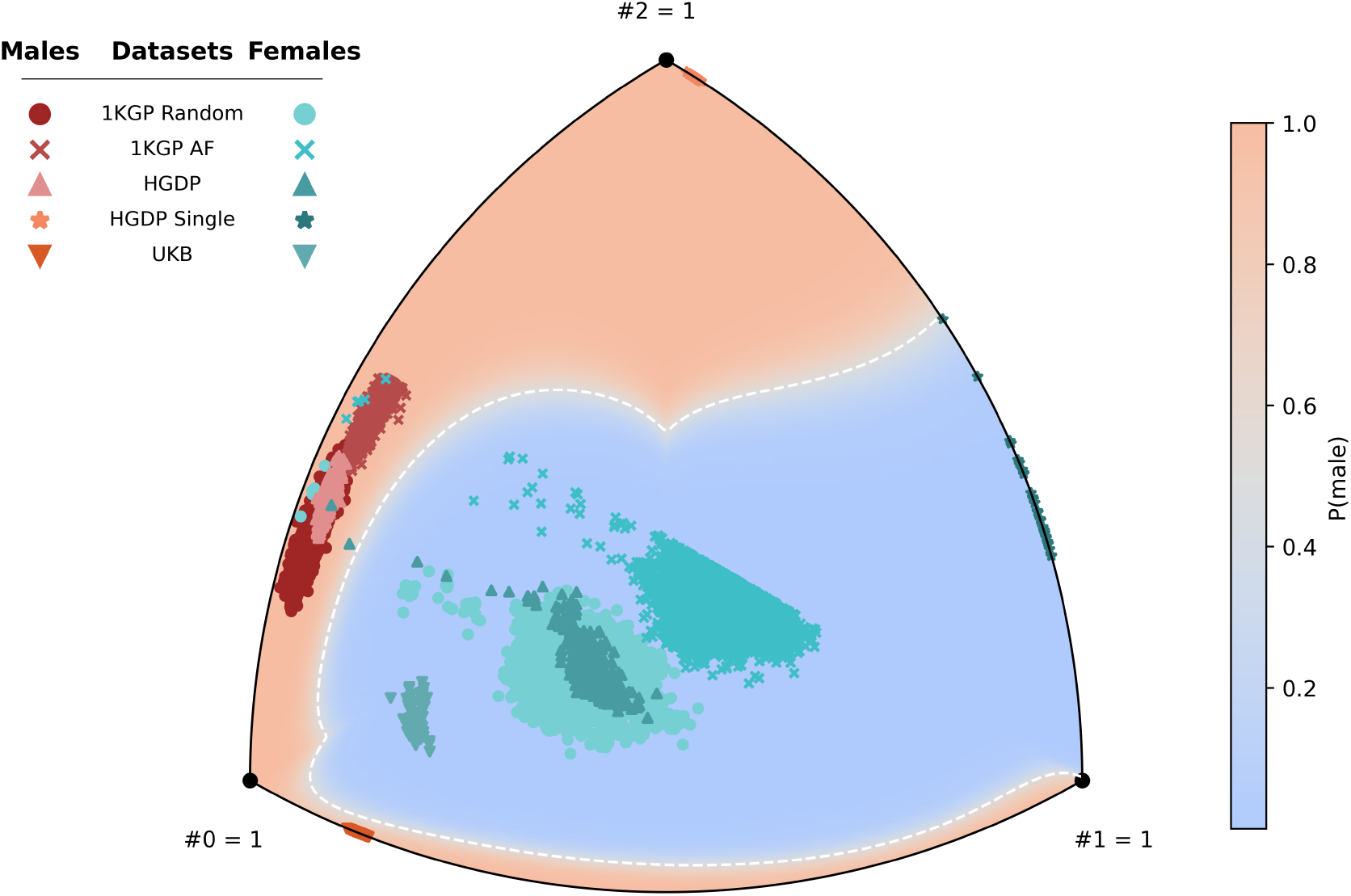
Universal sex-inference decision boundary visualized on the genotype frequency simplex. The Reuleaux triangle projection represents the normalized feature space of X-chromosome genotype counts (*f*_#0_ + *f*_#1_ + *f*_#2_ = 1). The background heatmap displays the learned probability surface, ranging from *P* (Male) ≈ 0 (Blue/Female) to *P* (Male) ≈ 1 (Orange/Male), with the white dashed line marking the decision threshold (*P* = 0.5). Overlaid points represent real samples from diverse benchmarks: WGS (1KGP, HGDP) whose males cluster is located between the #0 and #2 vertices; Arrays (UK Biobank) whose males are confined into the #0-#1 axis; and Single-Sample (HGDP Single) clustering near the #1-#2 axis. A single mathematical equation successfully classifies all modalities.

## 2 Methods

Zigo is a simple and interpretable sex checking method capable of generalizing across genomic technologies such as WGS, genotyping arrays, and single-sample extractions without relying on manual threshold tuning and reference panel homogenization. Additionally, Zigo is designed to be robust to any minor allele frequency (MAF) filtering imposed, and trained to classify samples correctly regardless of the ancestry. To achieve this, we first characterized the variability of genotype class distributions across different data modalities and subsequently developed a simulation-based training pipeline to capture this variance.

### 2.1 Zygosity Distributions

We analyzed X-chromosome genotype counts across multiple public datasets to assess the consistency of zygosity distributions. As illustrated in **Supplementary Figure S1**, the relative proportions of homozygous reference (0/0), heterozygous (0/1), and homozygous alternate (1/1) calls fluctuate significantly depending on the sequencing technology, MAF filtering, and population structure.

Males are biologically hemizygous in the non-pseudoautosomal region (non-PAR) of the X chromosome. However, standard variant calling pipelines typically represent these haploid calls as diploid homozygous genotypes (e.g., 0/0 or 1/1) to maintain VCF format consistency. Consequently, male distributions appear ostensibly diploid, characterized by a distinct absence of heterozygous calls (0/1). Notably, the genotype array sequences of the UK Biobank, as well as other array-based large-scale databases, enforces a strict haploid model for males, reporting true allelic counts (0 and 1). Furthermore, when performing single-sample sequencing and variant calling, many variant calling software will produce files with the reference homozygous calls (0/0) missing providing a zygosity distribution completely different than joint-calling WGS files and genotype-array files.

Given this heterogeneity, training a model on any single real dataset would likely result in poor generalization to other cases.

### 2.2 Synthetic Data Simulation

#### 2.2.1 Founder Population Simulation

We generated synthetic whole-genome variant calls using the `stdpopsim` library [2, 16], which serves as a standardized interface for the `msprime` coalescent simulation engine [14]. To ensure biological realism, we employed the ‘OutOfAfrica_3G09’ demographic model [12], which simulates the expansion of human populations into African (YRI), European (CEU), and East Asian (CHB) lineages. This model incorporates appropriate effective population sizes and generation times to produce realistic allele frequency spectra [13, 22]. Simulations were performed specifically for the X chromosome generating 1,000 samples per population, using the empirical recombination map from HapMap Phase II (GRCh38) [24] to accurately recapitulate linkage disequilibrium blocks and genetic architecture.

#### 2.2.2 Genetic Sex Modeling

Biological sex was assigned randomly to each simulated sample with equal probability *P* (Male) = *P* (Female) = 0.5. Since the coalescent engine outputs diploid genotypes by default (*G*_*in*_ = {*a*_1_, *a*_2_} where *a*_*i*_ ∈ {0, 1}), we applied a transformation function to model the specific chromosomal architecture of each sex and to inject realistic genotyping errors.

Let *g*_*ϵ*_ be the genotyping error rate, and let *δ* ∼ Bernoulli(*g*_*ϵ*_) be a random variable representing the occurrence of an error (bit-flip). We denote the bitwise XOR operation as ⊕, such that *x* ⊕ 1 = 1 −*x* (error) and *x*⊕ 0 = *x* (no error).

For **Females (XX)**, we modeled the genotype as fully diploid, treating both alleles independently. Genotyping errors were injected separately into each allele to simulate random sequencing or calling artifacts: *G*_female_ = {*a*_1_ ⊕ *δ*_1_, *a*_2_ ⊕ *δ*_2_ }, where *δ*_1_ and *δ*_2_ are independent realizations of the error variable.

For **Males (XY)**, to strictly enforce biological hemizygosity while adhering to the pseudo-diploid standard of VCF files, we discarded the second simulated allele (*a*_2_). The first allele (*a*_1_) was duplicated to fill the second slot. An error term was applied only to this duplicated allele to simulate the possibility of spurious heterozygous calls arising from technical noise: *G*_male_ ={ *a*_1_, *a*_1_ ⊕ *δ* }. This modeling ensures that male samples exhibit perfect homozygosity (0/0 or 1/1) in the absence of errors (*δ* = 0), but retain a non-zero probability of appearing heterozygous (0/1) proportional to *g*_*ϵ*_, reflecting real-world data imperfections.

#### 2.2.3 Allele Frequency Spectrum Augmentation

Standard genomic studies often apply disparate MAF filters during quality control, or target specific frequency ranges by design (e.g., rare variant associations vs. common variant arrays). To ensure our model remains robust regardless of the underlying frequency spectrum, we implemented a data augmentation step prior to technology emulation. We expanded the initial dataset of 3,000 simulated samples by generating 14 additional variations for each individual, applying a dense grid of empirically selected lower-bound MAF thresholds. These values were chosen to cover the full spectrum of possible frequency cutoffs, from rare variants to near-maximal entropy scenarios (detailed in Supplementary Note S1). Including the unfiltered baseline (MAF *>* 0), this process effectively multiplied the training set size by a factor of 15 (*N ≈* 45, 000 base profiles).

#### 2.2.4 Sequencing Technology and Format Emulation

To mimic distinct genotype encoding schemes—such as *post-hoc* haploid enforcement and the sparsity of single-sample VCFs—we generated three distinct representations for each simulated individual via deterministic transformations. Let **C** = [*c*_0_, *c*_1_, *c*_2_] be the vector of raw genotype counts for Homozygous Reference (0/0), Heterozygous (0/1), and Homozygous Alternate (1/1) calls, respectively, with normalized frequencies defined as **f** = **C**/ ∑**C**. From this baseline, we emulate three data formats:

1. **Joint-Call WGS (Baseline):** Standard population-scale VCFs where male hemizygosity is encoded as pseudo-diploid, using the raw output without modification: **C**_joint_ = **C**.
2. **Haploid-Encoded Array (Processed):** Processed biobank arrays with strict haploid encoding. Female vectors remain unchanged, but for **male samples**, the pseudo-diploid alternate class is merged into the haploid/heterozygous bin: 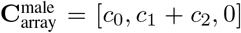. This forces the model to recognize males strictly adhering to a {0, 1} distribution, encapsulating both re-encoded alternate alleles and retained errors.
3. **Single-Sample:** VCFs where blocks of homozygous reference calls are omitted to conserve storage. We nullify the reference count, resulting in **C**_single_ = [0, *c*_1_, *c*_2_], with input features re-calculated as relative proportions: **f**_single_ = [0, *c*_1_/(*c*_1_ + *c*_2_), *c*_2_/(*c*_1_ + *c*_2_)]. This enables the model to detect sex-specific patterns solely from the ratio of non-reference calls. A pairwise visualization of the generated synthetic distributions is provided in Supplementary Figure S2.

### 2.3 Model Architecture and Knowledge Distillation

Our modeling strategy followed a two-stage framework designed to combine the high predictive power of gradient boosting decision trees (GBDT) with the portability and interpretability of a closed-form mathematical equation. Unlike SeXY, XYalign, and seGMM, Zigo operates without Y-chromosome or raw alignment data. Furthermore, unlike PLINK, Hail, peddy and BCFtools, it requires neither external reference panels nor manual threshold tuning (Supplementary Table S2).

#### 2.3.1 Gradient Boosting

We first trained a CatBoost [19] classifier to learn the decision boundaries between male and female genotype profiles. Cat-Boost was selected for its superior performance on tabular data and its robust handling of outliers. The model was trained on the augmented synthetic dataset described above, using the normalized genotype counts **x** = [*f*_0_, *f*_1_, *f*_2_] as input features. Hyperparameters were optimized using the Optuna automatic hyperparameter optimization framework [3] (Supplementary Table S1).

#### 2.3.2 Polynomial Distillation

While accurate, GBDT models require software dependencies, model storage, among other technical debt. To create a lightweight, dependency-free tool, we employed a knowledge distillation approach to approximate the decision boundary of the CatBoost model with a high-order polynomial equation.

We generated a dense grid of synthetic query points over the probability 2D-simplex (the space of all possible valid genotype frequencies) and queried the trained CatBoost model to obtain the predicted probability of being male, *p*_gbm_(**x**). To linearize the problem for regression, we transformed these probabilities into log-odds (logits) space:

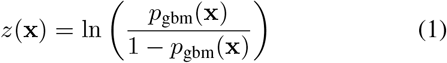

Probabilities were clipped to the range [*ϵ*, 1 − *ϵ*] prior to transformation in order to avoid numerical instability,

We then fitted a Polynomial Ridge Regression of degree *d* = 6 and regularization alpha *α* = 0.01 to these logits. The distilled model approximates the log-odds *z*(**x**) as a linear combination of polynomial basis functions *ϕ*(**x**):

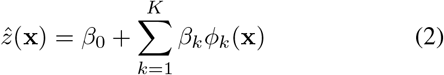

where *β* are the learned coefficients and *ϕ*_*k*_(**x**) represents the interaction terms of the input frequencies up to degree 6 (e.g., 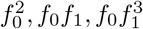, etc.). The degree was selected empirically to balance model complexity with approximation fidelity.

#### 2.3.3 Final Inference Equation

The final sex-checking tool operates by computing the polynomial score 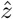 and converting it back to a probability space using the logistic sigmoid function:

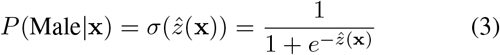

This formulation allows the entire inference logic to be encapsulated in a single mathematical expression comprising the polynomial coefficients, enabling instant classification with zero software overhead.

### 2.4 Evaluation Datasets and Benchmarking

To benchmark Zigo against established tools, we curated three distinct testing cohorts representative of the primary genomic data modalities.

#### 2.4.1 Data Sources

We obtained high-coverage Whole Genome Sequencing genotype calls for the 1000 Genomes Project (1KGP) and the Human Genome Diversity Project (HGDP) [1, 8] from the gnomAD v3.1 data portal. These datasets provide a diverse global representation, critical for assessing ancestry-agnostic performance. Ground-truth sex labels and metadata were sourced from the official 1000 Genomes FTP site (integrated call set v3.20200731) and the International Genome Sample Resource (IGSR) portal for HGDP.

To evaluate performance on array-based data structures, we utilized a random subset of 3,000 samples from the UK Biobank (UKB) [7]. As discussed in Section 2.1, this dataset serves as the standard for haploid-encoded male genotypes.

#### 2.4.2 Experimental Setup

To ensure a rigorous and fair benchmarking against established tools, particularly those that can leverage external population statistics (e.g., PLINK and Hail), we designed specific data partitions and stress-test scenarios.

##### Reference Panel Generation (1KGP Split)

Standard sex-checking methods often require accurate population allele frequencies to compute expected homozygosity rates. To provide these “priors” without data leakage, we partitioned the 1000 Genomes Project dataset (*N* = 3, 199) into two subsets:

- **Evaluation Set (***N* = 1, 600): Used exclusively for testing the performance of all models.
- **Reference Frequency Set (***N* = 1, 599): Used solely to calculate empirical allele frequencies 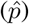. These calculated frequencies were provided as external input to PLINK and Hail during the benchmarking phase.

The 1KGP was selected for this purpose due to its high SNP density and global diversity, offering a more robust frequency spectrum than array-based datasets.

##### Single-Sample Emulation (HGDP-Single)

Since public single-sample datasets with reliable ground truth are scarce, we generated a single-sample dataset by extracting 100 random samples from the HGDP joint-called VCF into individual files. We removed all homozygous reference blocks (0/0) and strictly retained only sites with variation, effectively recreating the sparsity and distributional characteristics of isolated clinical samples obtained with commonly used variant calling pipelines. While this synthetic ablation effectively mimics the structural missingness of single-sample VCFs, we acknowledge that it may not capture every complex technical bias inherent to genuinely isolated single-called sequencing pipelines.

##### Robustness Stress-Testing

We assessed model stability under two distinct dimensions of data scarcity: variant count reduction and ascertainment bias.

###### 1. Random Downsampling (Variant Scarcity)

Starting from the full 1KGP evaluation set (*S*_0_, containing *M* ≈ 262, 991 variants), we generated a sequence of 10 subsets {*S*_1_, …, *S*_10_} by recursively random-subsampling half of the variants from the previous set:

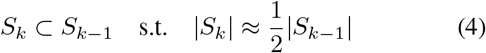

This yielded a testing gradient ranging from dense WGS down to extreme sparsity (| *S*_10_| = 513 variants).

###### 2. High-AF Filtering (Ascertainment Bias)

To evaluate robustness against datasets enriched for common variants (e.g., specific arrays or filtered sets), we generated a sequence of 5 subsets {*A*_1_, …, *A*_5_} by recursively retaining the top 50% of variants with the highest MAF.

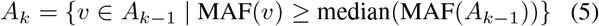

This process progressively increased the minimum AF from 0.25 (*A*_1_) to 0.48 (*A*_5_), simulating environments dominated exclusively by highly common polymorphisms.

## 3 Results

### 3.1 Geometric Separation of Sexes in the Genotype Simplex

To visually assess the discriminative power of our derived polynomial equation, we projected the normalized genotype class counts of the test datasets onto the 2D simplex visualized in Figure 1. We observed that real-world samples, despite originating from different sequencing technologies and processing pipelines, map to distinct but coherent regions within this geometric space. WGS male samples (1KGP, HGDP) cluster along the edge connecting the homozygous reference (#0) and alternate (#2) vertices, reflecting the pseudo-diploid encoding of males. Conversely, array-based male samples (UK Biobank) and single-sample extracts for both genders (HGDP-Single) cluster along the #0-#1 and #1-#2 axes, respectively, due to haploid encoding and reference block omission. Our model’s learned probability surface and decision boundary (white dashed line) successfully segregate male and female clusters across all distinct modalities simultaneously. Namely, the simplex visualization fully characterizes the model’s behavior, showing that it can accurately classify sex based on chromosome’s X zygosity.

### 3.2 General Performance

We benchmarked the performance of the proposed Zigo against three established tools: PLINK (v1.9), Hail, and BCFtools. Evaluations were performed on the 1000 Genomes Project (WGS), HGDP (WGS), and UK Biobank (Array) datasets. For PLINK and Hail, we assessed performance both with their default internal frequency estimation and with external population priors computed from a reference (w/ train), while BCFtools and Zigo were run in their standard modes.

As summarized in Table 1, our method achieved state-of-the-art accuracy across all cohorts, matching the performance of the probabilistic HMM-based method (BCFtools) and outperforming standard heterozygosity-based approaches. In the WGS cohorts (HGDP and 1KGP), Zigo yielded near-perfect classification (Balanced Accuracy *>* 0.999, ≤2 errors). In contrast, PLINK and Hail showed significantly higher error rates when using default thresholds, misclassifying between 44 and 220 samples depending on the dataset. While providing external frequencies improved the stability of PLINK and Hail, they did not surpass the accuracy of our reference-free approach. In the UK Biobank (Array) dataset, Zigo achieved 100% accuracy (0 errors), successfully handling the haploid-encoded males. Notably, `hail-f-train` exhibited a sharp performance drop (677 errors), likely due to mismatches between the WGS-derived priors and the array-specific allele frequencies.

**Table 1:** Benchmarking of sex inference performance across diverse genomic datasets. Comparison of Zigo against established tools (Hail, PLINK, BCFtools) across WGS, Array, and Single-Sample modalities. Methods were evaluated in their default modes and, where applicable, using external population priors (w/ train). **Min.#Err** indicates theoretical minimum errors with optimized post-hoc thresholds. Zigo achieves state- of-the-art accuracy across all scenarios without external priors.

| Method | Bal. Acc. | AUC | #Err | Min.#Err |
| --- | --- | --- | --- | --- |
| *Dataset: HGDP (WGS)* |  |  |  |  |
| Hail | 0.9051 | 0.9982 | 52 | 1 |
| Hail (w/ train) | 0.8668 | 0.9982 | 73 | 1 |
| PLINK | 0.9197 | 0.9999 | 44 | 1 |
| PLINK (w/ train) | 0.8923 | 0.9999 | 59 | 1 |
| **BCFtools** | 0.9964 | 0.9981 | **2** | 1 |
| **Zigo** | 0.9964 | 0.9999 | **2** | 1 |
| *Dataset: 1000 Genomes (WGS)* |  |  |  |  |
| Hail | 0.8647 | 1.0000 | 217 | 1 |
| Hail (w/ train) | 0.8628 | 1.0000 | 220 | 1 |
| PLINK | 0.8834 | 0.9988 | 187 | 1 |
| PLINK (w/ train) | 0.8747 | 0.9988 | 201 | 1 |
| **BCFtools** | 0.9994 | 0.9988 | **1** | 1 |
| **Zigo** | 0.9994 | 0.9988 | **1** | 1 |
| *Dataset: UKB (Array)* |  |  |  |  |
| Hail | 0.9947 | 1.0000 | 17 | 0 |
| Hail (w/ train) | 0.7892 | 0.7892 | 677 | 677 |
| PLINK | 0.9947 | 1.0000 | 17 | 0 |
| PLINK (w/ train) | 0.9885 | 1.0000 | 37 | 0 |
| **BCFtools** | 1.0000 | 1.0000 | **0** | 0 |
| **Zigo** | 1.0000 | 1.0000 | **0** | 0 |
| *Dataset: HGDP-Single (Single-Sample)* |  |  |  |  |
| Hail | 0.5000 | 0.5000 | 67 | 33 |
| Hail (w/ train) | 1.0000 | 1.0000 | 0 | 0 |
| PLINK | 0.5000 | 0.5000 | 67 | 33 |
| PLINK (w/ train) | 1.0000 | 1.0000 | 0 | 0 |
| BCFtools | 0.5000 | 1.0000 | 67 | 0 |
| **Zigo** | 1.0000 | 1.0000 | **0** | 0 |

Zigo significantly improves time efficiency. In our benchmarks on the HGDP dataset, Zigo required only 24.48 seconds for inference, matching the speed of PLINK (24.75s) and drastically outperforming both Hail (44.26s) and BCFtools (56.02s).

Results for Zigo presented in Table 1 reflect the distilled polynomial regression version (Supplementary Note S3). Comparative results for the CatBoost model are provided in Supplementary Fig. S3.

### 3.3 Reference-Free Inference in Single-Sample Scenarios

A critical limitation of many sex-inference algorithms is the inability to estimate allele frequencies internally during single-sample analysis, where the sample size is *N* = 1. We evaluated this scenario using the HGDP-Single dataset (Table 1).

In this regime, methods relying on internal statistics failed catastrophically. Hail and PLINK yielded a balanced accuracy of 0.5 (random guessing), as the lack of cohort-level data prevents calculating accurate expected homozygosity. Similarly, BCFtools, which defaults to estimating frequencies from the input stream, also failed to classify samples correctly (Balanced Accuracy ≈ 0.5) because the input provided insufficient information for its HMM priors. While PLINK and Hail achieved perfect accuracy when provided with external reference panels (w/ train), this dependency creates a bottleneck for clinical pipelines or isolated processing. These results highlight the distinct advantage of our approach over existing probabilistic tools: Zigo achieved perfect classification without requiring any external files or parameters, demonstrating its capability to handle sparse, isolated data by leveraging the learned geometric invariants of genotype counts.

### 3.4 Robustness to Variant Sparsity and Ascertainment Bias

Finally, we stress-tested the stability of our model under conditions of extreme data scarcity and ascertainment bias using the 1KGP robustness subsets (detailed results and methodology are provided in Supplementary Note S4).

While the accuracy of PLINK and Hail degraded at lower variant counts (resulting in a balanced accuracy of ∼ 0.875), Zigo maintained a flat error profile (0 errors) and perfect AUC throughout the entire gradient, matching the stability of BCFtools. Second, as the minimum MAF threshold increased, the balanced accuracy of PLINK and Hail fluctuated reflecting the sensitivity of the F-statistic to deviations from expected Hardy-Weinberg proportions in high-frequency regimes. Zigo remained strictly stable (100% accuracy). These results confirm that our simulation-based training strategy, which included spectrum augmentation, successfully conferred robustness against the distributional shifts caused by heavy variant filtering.

### 3.5 Discrepant Samples and Aneuploidy Detection

Despite the near-perfect accuracy of our method, a small number of samples were consistently classified as Male despite being labeled as Female in the ground-truth metadata. We investigated these discrepancies to determine if they represented algorithmic failure or genuine biological anomalies.

In the 1000 Genomes Project, the sample *HG03511* is labeled as female in the official pedigree file but was classified as male by Zigo (and BCFtools). According to the PLINK 2.0 resource documentation, detailed quality control analysis reveals that this cell line exhibits coverage and heterozygosity statistics consistent with a single copy of chromosome X and no Y chromosome (karyotype 45,X or mosaic) [9]. While labeled female, the sample is genetically hemizygous for the X chromosome. Therefore, our model correctly identified the underlying haploid genetic architecture, effectively flagging a case of likely Turner syndrome or mosaic loss of chrX.

Similarly, in the HGDP dataset, two samples labeled as female—*HGDP01239* (Daur) and *HGDP01273* (Mozabite)— were classified as male. Inspection of their genotype counts reveals a complete absence of heterozygous calls despite high coverage, placing them firmly in the male/hemizygous cluster of the simplex (Figure 1). These results strongly suggest that these samples also harbor unreported X-chromosome monosomies or severe mosaicism indistinguishable from hemizygosity in genotype space. These findings highlight the utility of Zigo as a sensitive QC tool for detecting sex-chromosomal aneuploidies (e.g., X0 females) that standard pedigree checks might miss.

## 4 Discussion

Our reference-free method, Zigo, infers genetic sex from X-chromosome genotype data with state-of-the-art accuracy across diverse sequencing technologies, robustly handling severe data sparsity and isolated single-samples where traditional internal frequency estimation fails. By distilling a simulation-trained gradient boosting model into a single polynomial equation, we provide a tool that is both mathematically interpretable and computationally lightweight, eliminating the need for external reference panels, Y-chromosome data, or manual tuning.

A key insight is that the geometric distribution of normalized genotype counts contains sufficient information to discriminate sex, transcending specific file formats. However, standard F-statistic methods assume a relationship between observed and expected homozygosity that breaks down when allele frequencies are unknown or biased by array design [6]. While these methods perform adequately given accurate external priors, their performance degrades significantly in isolation. In contrast, Zigo leverages invariant geometric signatures of sex-specific zygosity consistent across major biobanks and diverse ancestral populations, allowing seamless generalization even to single-sample VCFs where reference block omission catastrophically biases standard metrics.

Explicitly modeling both *pseudo-diploid* VCF encodings and *haploid* array encodings prevented overfitting to specific standards. Stress-tests proved this strategy’s utility, maintaining perfect accuracy even with under 600 variants or restricted common polymorphisms. This resilience makes Zigo ideal for targeted panels or low-pass sequencing where sparsity breaks other methods. Additionally, distilling the model into a dependency-free, closed-form equation facilitates seamless pipeline integration and reproducibility.

A notable limitation is that, despite simulating broad demographic histories [12], we did not explicitly model complex aneuploidies such as XXY (Klinefelter syndrome) or X0 (Turner syndrome). However, our empirical detection of likely X0 samples indicates that these anomalies naturally map to the hemizygous region of the genotype simplex, effectively flagging them as discordant. To fully characterize the geometric signatures of broader aneuploidies and quantify the method’s diagnostic sensitivity, dedicated validation on clinical cytogenetic datasets is warranted.

In conclusion, as genomic datasets continue to grow in scale and heterogeneity, QC tools must evolve to be as robust and automated as the analysis pipelines they support. We anticipate that reference-free, simulation-trained methods like Zigo will become standard for harmonizing quality control across decentralized and privacy-sensitive data environments, where sharing individual-level reference data is not feasible.

## 5 Acknowledgments

We thank Cole Shanks for valuable feedback and Ferran Marqués for his initial supervision. This work uses data provided by patients and collected by the NHS as part of their care and support. We thank the UK Biobank and its participants and the NHS. This research has been conducted using the UK Biobank Resource under Application Number 24983.

## Supplementary Material

### 1. Supplementary Notes

#### Supplementary Note S1: Allele Frequency Spectrum Augmentation Thresholds

During the data augmentation step described in the main text, we expanded the simulated dataset by applying a grid of lower-bound Minor Allele Frequency (MAF) thresholds. Rather than using a strict linear or geometric progression, these specific values were selected empirically to densely and comprehensively cover the feasible range of frequency cutoffs encountered in diverse genomic studies (from rare variant analyses to arrays enriched for common variants).

The 14 specific thresholds applied are:

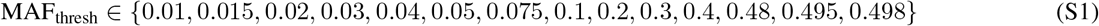

#### Supplementary Note S2: CatBoost Model Configuration

To ensure the reproducibility of the experiments described in the main text, we detail the specific hyperparameters used to train the CatBoost classifier. These parameters remained fixed for all reported results and can be found in **Supplementary Table S1**.

#### Supplementary Note S3: Explicit Formulation of the Distilled Polynomial Regression

The distilled version of the Zigo method utilizes a polynomial regression model of degree 6 to approximate the decision boundary of the original ensemble. The probability of the target class (male), denoted as *P* (*y* = 1|**x**), is modeled using a logistic function:

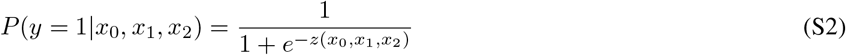

where *z*(**x**) is the logit function defined as a polynomial expansion of the input features *x*_0_, *x*_1_, and *x*_2_. The general form is:

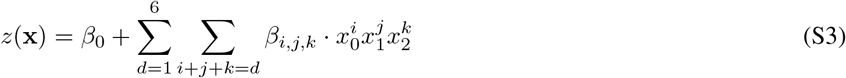

The specific coefficients (*β*) derived from the distillation process are detailed in **Supplementary Table S3**. To assess the efficacy of knowledge distillation, we compared the Distilled Polynomial Regression against both the original CatBoost and a baseline Polynomial Regression trained directly on the raw data (**Supplementary Table S4**). The distilled version bridges the performance gap, significantly outperforming the baseline and matching the gradient boosting predictive power.

#### Supplementary Note S4: Detailed Robustness Stress-Testing Evaluation

To assess model stability under extreme data degradation, we evaluated performance along the two generated dimensions (**Supplementary Figure S4**).

For variant scarcity, the number of available SNPs was recursively halved. We observed that methods based on heterozygosity ratios (PLINK, Hail) experience a significant decrease in classification performance when provided with fewer than 1,027 variants, pointing to a minimum SNP threshold required for medium-quality results. Conversely, Zigo and BCFtools maintained perfect stability. For ascertainment bias, progressively higher Minor Allele Frequency (MAF) thresholds were applied to simulate arrays enriched exclusively for common variants. While F-statistic methods showed sensitivity to deviations from expected Hardy-Weinberg proportions, Zigo maintained perfect classification.

### 2. Supplementary Figures

**Figure S1:**
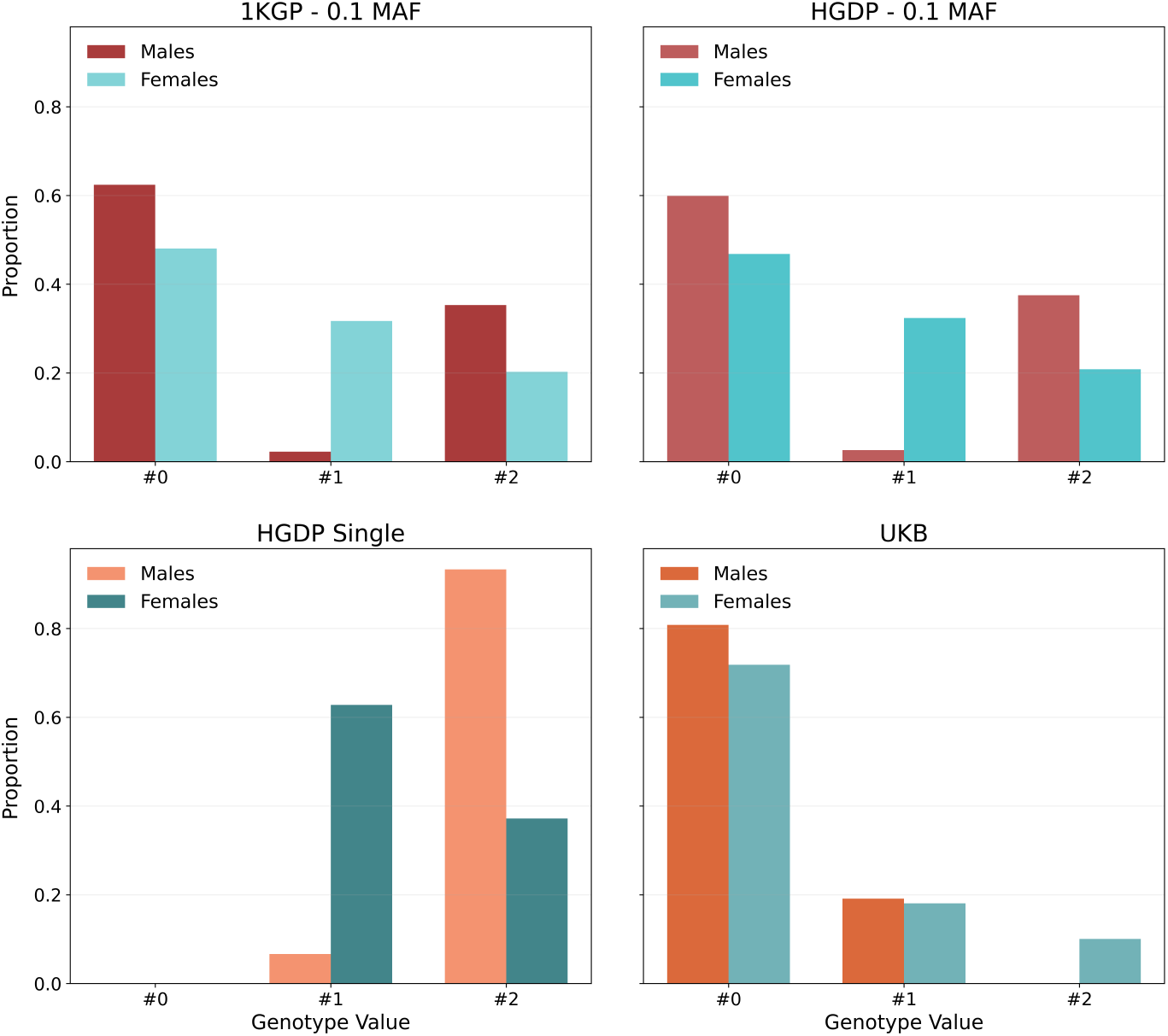
Variability of X-chromosome genotype distributions across real datasets. The bar plots show the normalized proportions of Reference (#0), Heterozygous (#1), and Alternate (#2) genotype calls for males (red/orange) and females (blue/teal) across distinct datasets (1000 Genomes, HGDP, UK Biobank). Significant discrepancies are observed between sequencing technologies (WGS vs. Array). HGDP Single (a subset of 100 HGDP samples) illustrates the distributional shift inherent to single-sample formats, where the absence of homozygous reference calls drastically alters the frequency spectrum compared to the full joint-called HGDP dataset. These variations highlight the need for a training strategy that encompasses this heterogeneity.

**Figure S2:**
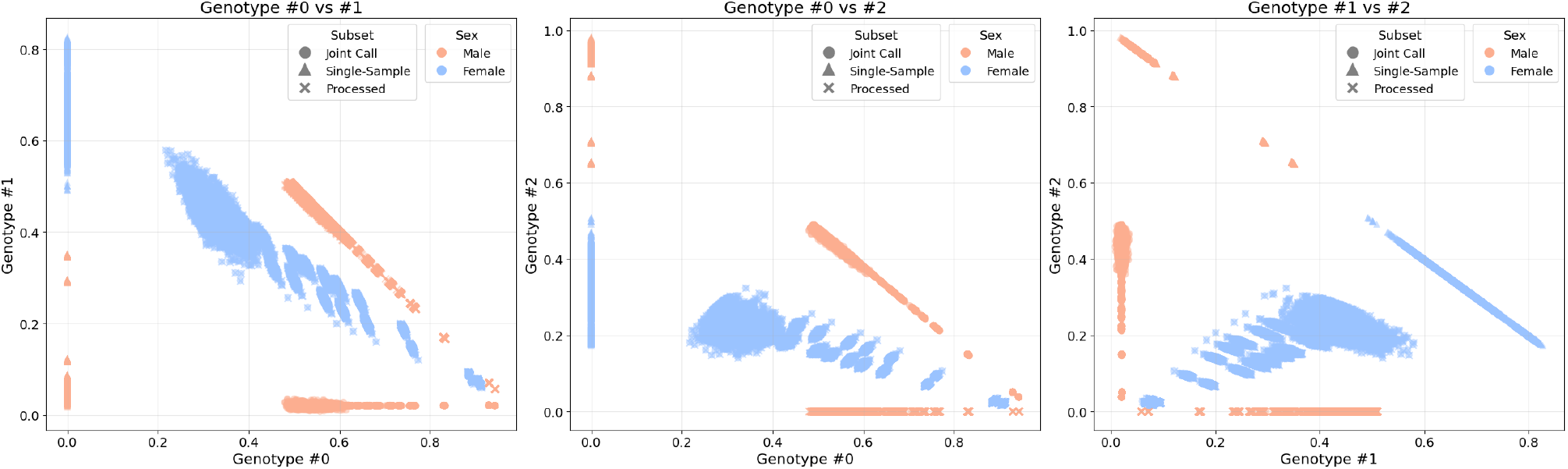
Synthetic genotype frequency landscapes. Pairwise projections of normalized genotype class counts (#0 Ref, #1 Het, #2 Alt) for the augmented training dataset. The scatter plots illustrate the distinct distributional signatures generated for Joint-Called (circles), Single-Sample (triangles), and Processed/Array (crosses) formats across biological sexes (Males in orange, Females in blue). Note how *Processed* males are strictly confined to the axis where Genotype #2 is zero (reflecting haploid encoding), while *Single-Sample* profiles cluster at Genotype #0 are zero (reflecting the omission of reference blocks).

**Figure S3:**
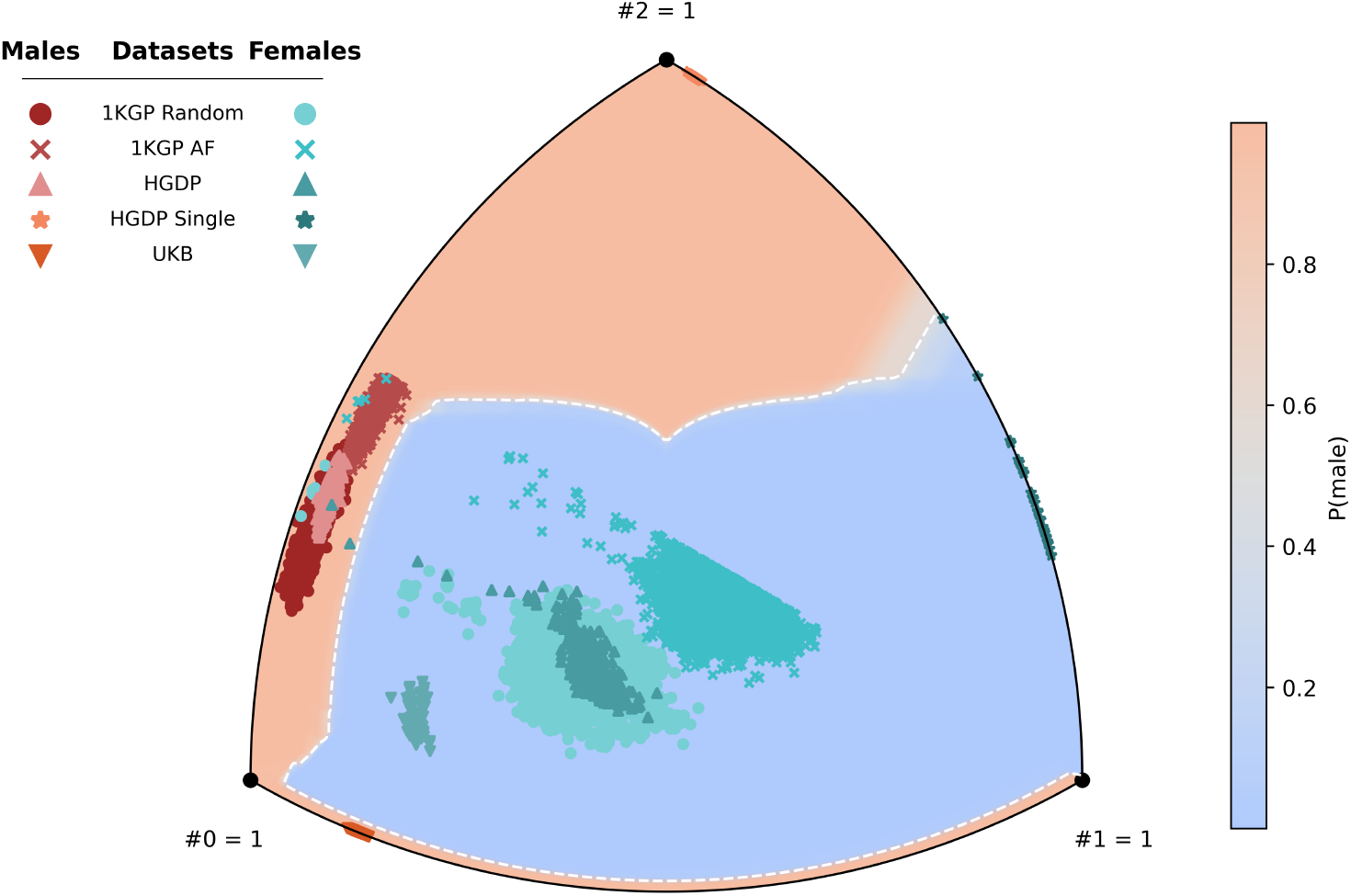
Performance of Zigo’s Catboost-based model. Decision boundary learned by the original gradient boosting model projected onto the Reuleaux triangle representation of the genotype frequency simplex. This illustrates the complexity of the original decision boundary prior to polynomial distillation.

**Figure S4:**
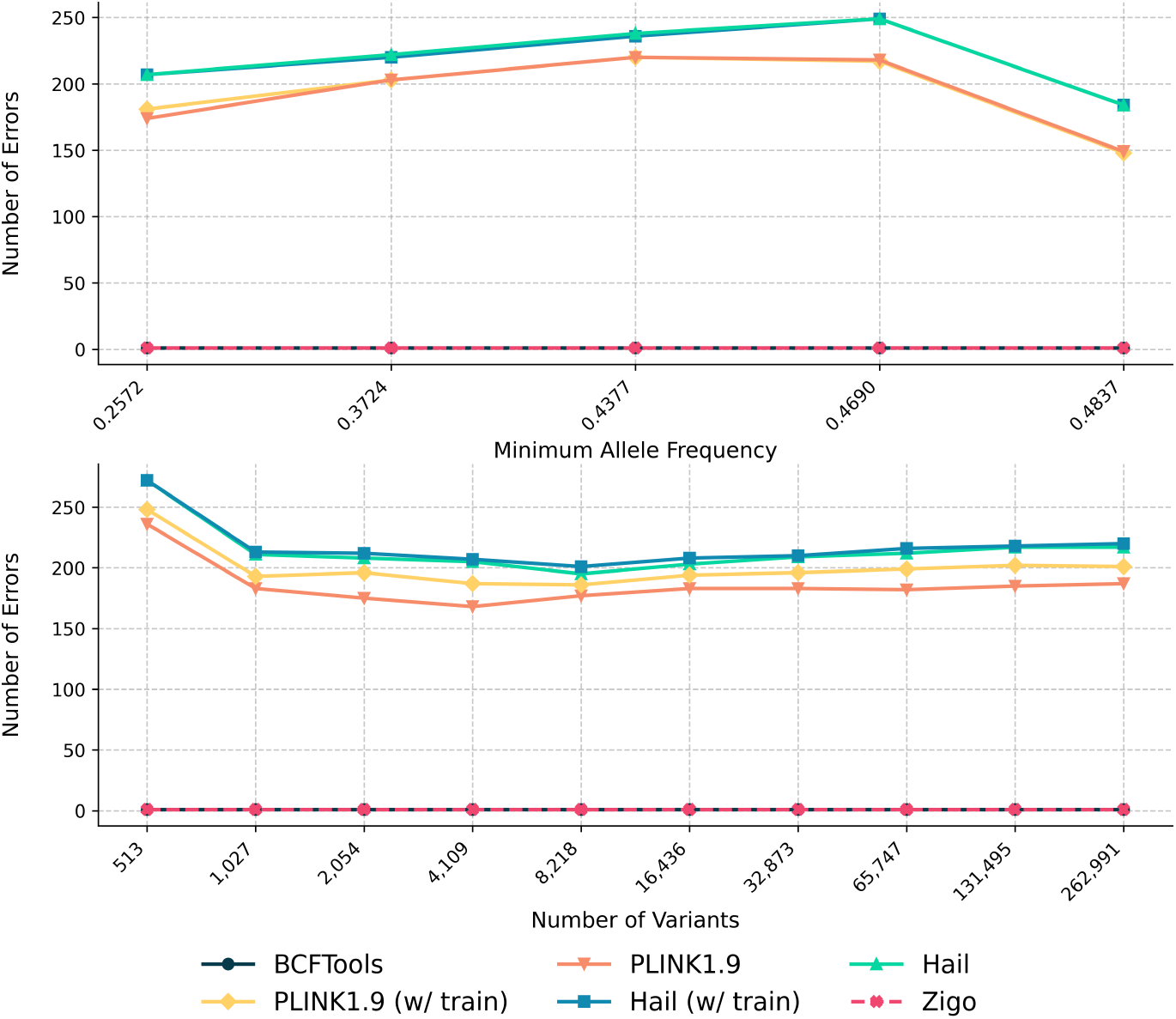
Robustness stress-testing under data scarcity and ascertainment bias. Number of Errors on the 1000 Genomes Evaluation set under progressive degradation. **Ascertainment Bias (Top):** Performance as the MAF threshold is increased from 0.25 to 0.48, simulating datasets enriched for common variants. **Variant Scarcity (Bottom):** Performance as the total number of variants is recursively halved from ∼263,000 to 513. While methods based on heterozygosity ratios show sensitivity to high MAF and degradation at low variant counts, Zigo maintains perfect stability across all regimes, matching the robustness of BCFtools.

### 3. Supplementary Tables

**Table S1:** CatBoost hyperparameters. Hyperparameters used for the CatBoost model training.

| Parameter | Value |
| --- | --- |
| Iterations | 978 |
| Depth | 6 |
| Learning Rate | 0.0595 |
| L2 Leaf Regularization | 9.3333 |
| Random Seed | 2003 |
| Loss Function | LogLoss |

**Table S2:** Operational capabilities of sex-inference methods. Autonomy of each tool based on four attributes: (1) Works without Y-chromosome data; (2) Operates without raw alignment files; (3) No need of external reference panels for frequency estimation in single-sample; (4) No manual tuning of decision thresholds required.

| Method | Y Free | Raw Free | Ref Free | Thr. Free |
| --- | --- | --- | --- | --- |
| SeXY | ✗ | ✗ | ✓ | ✗ |
| XYalign | ✗ | ✗ | ✓ | ✓ |
| seGMM | ✗ | ✗ | ✓ | ✓ |
| Hail | ✓ | ✓ | ✗ | ✗ |
| PLINK | ✓ | ✓ | ✗ | ✗ |
| peddy | ✓ | ✓ | ✗ | ✗ |
| BCFtools | ✓ | ✓ | ✗ | ✓ |
| **Zigo** | ✓ | ✓ | ✓ | ✓ |

**Table S3:** Coefficients for the Degree-6 Polynomial Distilled Model. The terms are grouped by polynomial degree.

| Term | Coefficient | Term | Coefficient |
| --- | --- | --- | --- |
| *Degree 0 (Intercept)* |  |  |  |
| $\beta_0$ | -19.2882 | | |
| *Degree 1 (Linear)* |  |  |  |
| $x_0$ | -21.0116 | $x_1$ | 4.8968 |
| $x_2$ | 33.5459 | | |
| *Degree 2 (Quadratic)* |  |  |  |
| $x_0^2$ | -33.8280 | $x_0x_1$ | 25.3936 |
| $x_0x_2$ | -17.4267 | $x_1^2$ | 10.7148 |
| $x_1x_2$ | -37.9990 | $x_2^2$ | 77.7789 |
| *Degree 3 (Cubic)* |  |  |  |
| $x_0^3$ | 42.2327 | $x_0^2x_1$ | -12.4375 |
| $x_0^2x_2$ | -54.7341 | $x_0x_1^2$ | 116.9304 |
| $x_0x_1x_2$ | -72.0918 | $x_0x_2^2$ | 107.6352 |
| $x_1^3$ | -86.3315 | $x_1^2x_2$ | -15.8324 |
| $x_1x_2^2$ | 50.1615 | $x_2^3$ | -82.6634 |
| *Degree 4* |  |  |  |
| $x_0^4$ | 80.2211 | $x_0^3x_1$ | -57.0134 |
| $x_0^3x_2$ | 16.4472 | $x_0^2x_1^2$ | 223.5260 |
| $x_0^2x_1x_2$ | -180.4350 | $x_0^2x_2^2$ | 109.2212 |
| $x_0x_1^3$ | -130.2908 | $x_0x_1^2x_2$ | 23.6009 |
| $x_0x_1x_2^2$ | 83.9444 | $x_0x_2^3$ | -79.9672 |
| $x_1^4$ | 80.5445 | $x_1^3x_2$ | -41.0813 |
| $x_1^2x_2^2$ | 0.9737 | $x_1x_2^3$ | -33.0964 |
| $x_2^4$ | 17.0367 | | |
| *Degree 5* |  |  |  |
| $x_0^5$ | 30.8818 | $x_0^4x_1$ | -20.5522 |
| $x_0^4x_2$ | 70.0352 | $x_0^3x_1^2$ | 205.5349 |
| $x_0^3x_1x_2$ | -242.2518 | $x_0^3x_2^2$ | 188.8652 |
| $x_0^2x_1^3$ | -24.7348 | $x_0^2x_1^2x_2$ | 42.4477 |
| $x_0^2x_1x_2^2$ | 19.0546 | $x_0^2x_2^3$ | -98.8647 |
| $x_0x_1^4$ | -8.8163 | $x_0x_1^3x_2$ | -95.8294 |
| $x_0x_1^2x_2^2$ | 76.9310 | $x_0x_1x_2^3$ | -11.8962 |
| $x_0x_2^4$ | 31.0151 | $x_1^5$ | 68.1550 |
| $x_1^4x_2$ | 22.4288 | $x_1^3x_2^2$ | 30.8480 |
| $x_1^2x_2^3$ | -106.4589 | $x_1x_2^4$ | 83.4009 |
| $x_2^5$ | -87.6807 | | |
| *Degree 6* |  |  |  |
| $x_0^6$ | -72.6799 | $x_0^5x_1$ | 44.9380 |
| $x_0^5x_2$ | 56.0432 | $x_0^4x_1^2$ | 188.7307 |
| $x_0^4x_1x_2$ | -252.9932 | $x_0^4x_2^2$ | 266.4874 |
| $x_0^3x_1^3$ | 15.5671 | $x_0^3x_1^2x_2$ | 1.4455 |
| $x_0^3x_1x_2^2$ | 9.5149 | $x_0^3x_2^3$ | -87.4453 |
| $x_0^2x_1^4$ | -31.3198 | $x_0^2x_1^3x_2$ | -9.0447 |
| $x_0^2x_1^2x_2^2$ | 50.1517 | $x_0^2x_1x_2^3$ | -40.6091 |
| $x_0^2x_2^4$ | 28.9439 | $x_0x_1^5$ | 156.9073 |
| $x_0x_1^4x_2$ | -134.7908 | $x_0x_1^3x_2^2$ | 47.9578 |
| $x_0x_1^2x_2^3$ | -21.1445 | $x_0x_1x_2^4$ | 49.6603 |
| $x_0x_2^5$ | -48.5155 | $x_1^6$ | -57.5880 |
| $x_1^5x_2$ | -32.3020 | $x_1^4x_2^2$ | 189.7865 |
| $x_1^3x_2^3$ | -206.6885 | $x_1^2x_2^4$ | 121.1713 |
| $x_1x_2^5$ | -87.7823 | $x_2^6$ | 69.4359 |

Note: Coefficients are rounded to 4 decimal places for display. Inputs $x_0, x_1, x_2$ correspond to $f_{\#0}, f_{\#1}, f_{\#2}$ and must be normalized ( $f_{\#0} + f_{\#1} + f_{\#2} = 1$ )

**Table S4:**
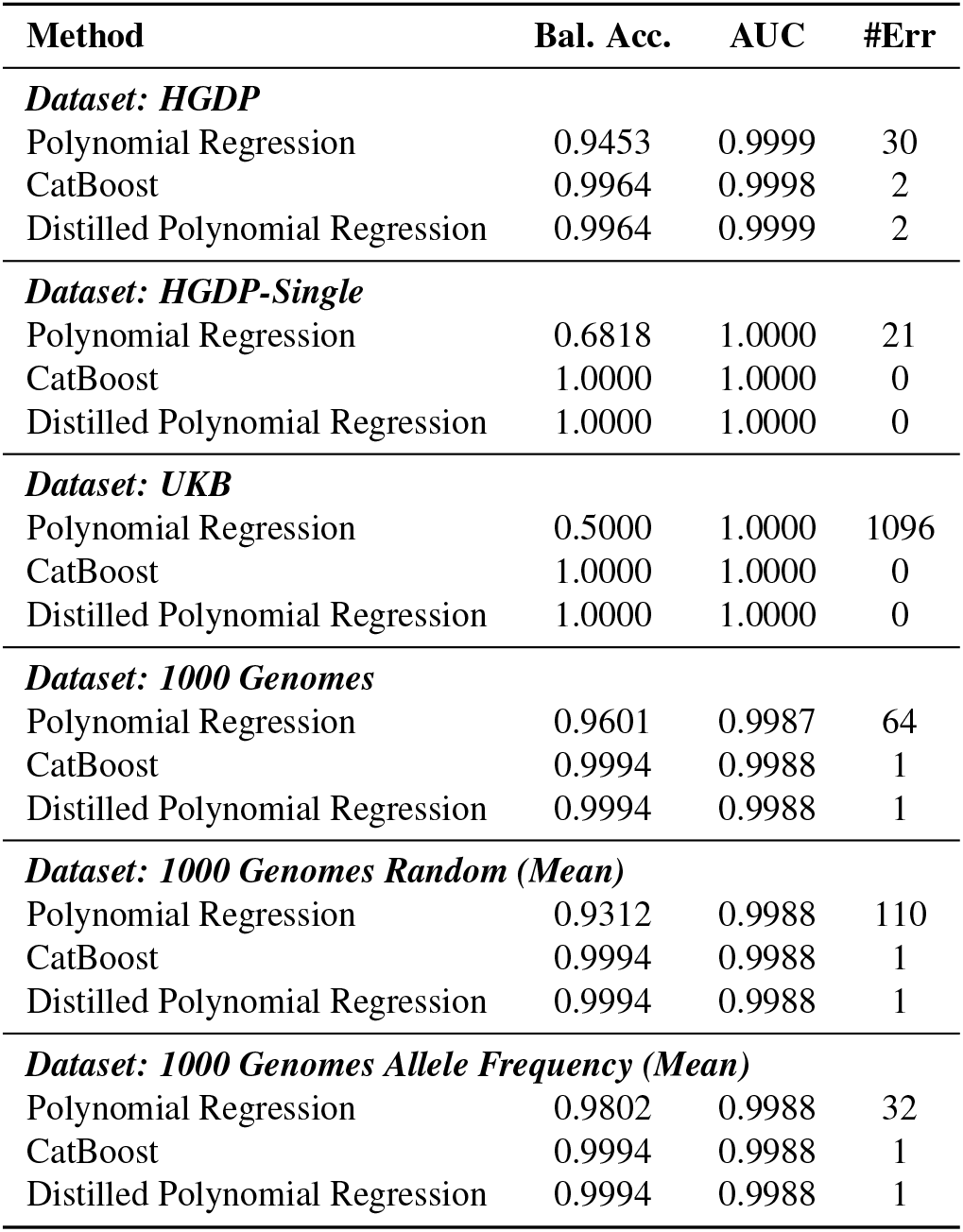
Performance comparison between the Baseline, Gradient Boosting and Distilled models. The evaluation metrics show that the lightweight polynomial regression maintains the robustness of the complex CatBoost ensemble while improving upon a direct baseline approach.

| Method | Bal. Acc. | AUC | #Err |
| --- | --- | --- | --- |
| ***Dataset: HGDP*** |  |  |  |
| Polynomial Regression | 0.9453 | 0.9999 | 30 |
| CatBoost | 0.9964 | 0.9998 | 2 |
| Distilled Polynomial Regression | 0.9964 | 0.9999 | 2 |
| ***Dataset: HGDP-Single*** |  |  |  |
| Polynomial Regression | 0.6818 | 1.0000 | 21 |
| CatBoost | 1.0000 | 1.0000 | 0 |
| Distilled Polynomial Regression | 1.0000 | 1.0000 | 0 |
| ***Dataset: UKB*** |  |  |  |
| Polynomial Regression | 0.5000 | 1.0000 | 1096 |
| CatBoost | 1.0000 | 1.0000 | 0 |
| Distilled Polynomial Regression | 1.0000 | 1.0000 | 0 |
| ***Dataset: 1000 Genomes*** |  |  |  |
| Polynomial Regression | 0.9601 | 0.9987 | 64 |
| CatBoost | 0.9994 | 0.9988 | 1 |
| Distilled Polynomial Regression | 0.9994 | 0.9988 | 1 |
| ***Dataset: 1000 Genomes Random (Mean)*** |  |  |  |
| Polynomial Regression | 0.9312 | 0.9988 | 110 |
| CatBoost | 0.9994 | 0.9988 | 1 |
| Distilled Polynomial Regression | 0.9994 | 0.9988 | 1 |
| ***Dataset: 1000 Genomes Allele Frequency (Mean)*** |  |  |  |
| Polynomial Regression | 0.9802 | 0.9988 | 32 |
| CatBoost | 0.9994 | 0.9988 | 1 |
| Distilled Polynomial Regression | 0.9994 | 0.9988 | 1 |

